# Diffusive dynamics of Aspartate *α*-decarboxylase (ADC) liganded with D-serine in aqueous solution

**DOI:** 10.1101/2020.08.11.244939

**Authors:** Tushar Raskar, Stephan Niebling, Juliette M. Devos, Briony A. Yorke, Michael Härtlein, Nils Huse, V. Trevor Forsyth, Tilo Seydel, Arwen R. Pearson

**Affiliations:** Institut Max von Laue - Paul Langevin, 71 Avenue des Martyrs, Grenoble 38000, France; Partnership for Structural Biology, 71 Avenue des Martyrs, Grenoble 38000, France; Institute for Nanostructure and Solid State Physics, Hamburg Centre for Ultrafast Imaging, Universität Hamburg, Luruper Chaussee 149, Hamburg, 22761, Germany; European Molecular Biological Laboratory, Hamburg outstation, Notkestraße 85, 22607 Hamburg, Germany; School of Chemistry and Bioscience, University of Bradford, Bradford, BD7 1DP, UK; Faculty of Natural Sciences, Keele University, Staffordshire, ST5 5BG, UK

## Abstract

Incoherent neutron spectroscopy, in combination with dynamic light scattering was used to investigate the effect of ligand binding on the center-of-mass self-diffusion and internal diffusive dynamics of *E.coli* aspartate *α*-decarboxylase (ADC). The X-ray crystal structure of the D-serine inhibitor complex with ADC was also determined, and molecular dynamics simulations used to further probe the structural rearrangements that occur as a result of ligand binding. These experiments reveal the existence of higher order oligomers of the ADC tetramer on ns-ms time-scales, and also show that ligand binding both affects the ADC internal diffusive dynamics and appears to further increase the size of the higher order oligomers.

## 1 Introduction

It is becoming increasingly clear that an understanding of the structure-function relationships of biological macromolecules, such as enzymes, requires both an understanding of their structure and of their dynamics. Enzymes, biological catalysts, are extremely large compared to chemical catalysts and are capable of very high specificity and selectivity, steering and controlling chemical reactions to usually produce a single desired outcome. Although the active site, where catalysis occurs, is usually compact and localised, the whole enzyme contributes to the enzymatic reaction [1, 2]. Biological macro-molecules can be considered as a soft elastic network and exhibit dynamics ranging over many orders in time, from femtosecond chemical steps to much slower millisecond to second large scale conformational rearrangements [3]. How these dynamics couple together to allow the slow motions of the protein to modulate catalysis remains an open question in structural enzymology and biophysics [4, 5]. There are multiple ways to probe these dynamics, including experiments that examine the macromolecule as it proceeds through its reaction cycle [6, 7], as well as methods that probe the equilibrium or non-driven dynamics [8]. However, many of the experimental tools to probe equilibrium dynamics require incorporation of labels, can access only dilute macromolecular suspensions, or cause radiation damage [9, 10]. Quasi-elastic neutron scattering (QENS) has been shown to be sensitive to the functionally relevant intra- and intermolecular dynamics of proteins [11, 12, 13], and how these dynamics change in response to ligand binding [14] or covalent modification [15]. These studies showed that binding of a substrate or its analogue can lead not only to local structural rearrangements in the binding site, but also to a larger change in the overall intra- and the intermolecular dynamics of the whole protein.

Here we study the *E. coli* enzyme aspartate *α*-decarboxylase (ADC), complementing previous work on off-the-shelf proteins in aqueous solution, [11] establishing a framework to systematically test the effect of parameter changes in proteins. In this case, this parameter is the presence or absence of the ligand.

ADC catalyzes the oxidative decarboxylation of L-aspartate to yield *β*-alanine [16]. *β*-alanine is required for the biosynthesis of pantothenate [17] which is then further converted to the important metabolic cofactor, Coenzyme A [16]. ADC contains a covalently linked protein-derived pyruvoyl cofactor which forms a Schiff base with the substrate [18] to initiate the decarboxylation reaction. This protein derived cofactor is formed via the posttranslational cleavage of the ADC zymogen protein backbone into *α* and *β* chains, resulting in the formation of a new C-terminus on the *α* chain and a N-terminal pyruvoyl cofactor on the *β* chain [19, 18]. The *α* C-terminus is extremely flexible, and initial binding of the substrate is associated with its rearrangement to close over the active site. This conformational rearrangement is believed to play a role in determining the overall rate of catalysis [20]. D-Serine is an inhibitor of ADC that, like the substrate L-aspartate binds in the enzyme active site [21].

## 2 Experiments and Methods

### 2.1 Sample preparation

ADC was expressed and purified according to previously published protocols [16, 18]. The final purified protein was concentrated using a 10 kDa cutoff centrifugal unit until a concentration of 135 mg/mL was reached. The buffer used for the final concentration was 50 mM Tris-HCl pH 7.4, 100 mM NaCl and 0.1 mM DTT in H_2_O. This was exchanged to a D_2_O buffer with the same composition for QENS measurements.

### 2.2 Crystallization of ADC and soaking with D-Serine

The protein was concentrated to 10 mg/mL and was crystallized by vapour diffusion after mixing the protein and the precipitant in 1:1 ratio. The best crystals were obtained in 1.8 M Ammonium sulphate, 100 mM Sodium citrate pH 4.5. D-Serine was dissolved in the crystallization buffer to a concentration of 1M before being added in a 1:1 ratio to the crystallisation droplet. The crystals were soaked for approximately 2 minutes before being washed in fresh crystallization buffer and then cryoprotected in crystallization buffer containing 20% *v/v* glycerol and flash cooled in liquid nitrogen.

### 2.3 X-ray data collection, processing and model refinement

X-ray diffraction data were collected at 100K at Diamond Light Source on beamline I24 using a Pilatus3 6M detector. Data were integrated, processed and scaled using XDS [22]. The structure was solved using the molecular replacement method as implemented in the software PHASER of the CCP4 suite [23]. The structure 1AW8 from the PDB was used as the search model after removing the ligands and the water molecules. Crystallographic refinement was carried out using REFMAC5 [24] from the CCP4 suite [23]. COOT [25] was used for real-space modelling.

### 2.4 Theoretical Diffusion Coefficient Calculation

The program HYDROPRO [26] was used to calculate the diffusion coefficients for the apo and D-serine bound ADC using the apo-ADC structure 1AW8 [19] and D-serine complex structure determined in this work as inputs. The partial specific volume of ADC was calculated as 0.70 cm^3^/g. The solvent viscosity for D_2_O was set to 0.01830, 0.01175, 0.00830 poise for the temperatures *T* = 280, 295 and 310 K, respectively [27].

### 2.5 Dynamic Light Scattering (DLS)

Dynamic light scattering experiments were conducted on an ALV-7004 instrument for both the ADC-APO and ADC-LIG samples. The recorded measurements covered the scattering angles from 30 to 150*°*. All measurements were conducted at *T* = 298.8 K. The ADC concentrations used covered a range from 1 to 5 mg/mL (16.67 to 83.3 *µ*M). The concentration of D-Serine used was 20 times that of the ADC tetramer (in terms of *µ*M) for each of the samples to ensure complete saturation of all the binding sites. The solvent used was D_2_O.

### 2.6 Protein concentration determination

Data were recorded using a DeNovix DS-11 spectrophotometer. Protein concentration was determined from absorbance at 280 nm and a molar extinction coefficient for ADC of. The accuracy of the spectrophotometric measurements was confirmed by dialysis against 20 mM ammonium acetate, pH 7.0, 100 mM NaCl, lyophilyzing and weighing a known amount of ADC. The measurements were carried out at an average temperature of *T* = 298.5 K. Lyophilization was carried out using a Martin Christ instrument at a vacuum pressure of 0.06 mbar.

### 2.7 Neutron spectroscopy experiments

Experiments were performed on solutions of ADC in D_2_O using both the IN16B and IN5 spectrometers at the ILL [28, 29]. IN16B [30, 31] has an energy resolution of 0.75 *µ*eV FWHM at 6.27 Å (Si111 crystal analyzer configuration), and IN5 has an energy resolution of approximately 80 *µ*eV FWHM at 5 Å incident wavelength. A cylindrical double walled aluminium sample holder sealed with indium wire was used for the measurements, with the difference in the radius between the two walls being 0.15 mm and the outer diameter 22 mm. The total liquid sample volume was 1.2 mL. The identical samples were used consecutively in both the IN16B and associated IN5 experiments. The temperature was controlled with a standard Orange cryostat.

### 2.8 Sample details for the neutron spectroscopy experiments

#### 2.8.1 Experiment 1

The concentration of ADC was 50 mg/mL corresponding to a dry protein volume fraction of 0.03 [32] and translating to 6.82 × 10^23^ hydrogens in total. The concentration of D-Serine was 30 mg/mL which translates to 12.046 × 10^23^ hydrogens in total.

#### 2.8.2 Experiment 2

The total quantity of ADC used was 162.3 mg, corresponding to a dry protein volume fraction of 0.09 [32]. To ensure the complete saturation of all the ADC binding sites, 45 mg of D-Serine was added per mL of the sample volume (135 mg/mL of ADC). This means there are a total of 18.29 × 10^23^ protons from the D-Serine against 18.07×10^23^ protons from the ADC molecule and that there are 47.6 molecules of D-Serine per ADC monomer. The protein solution and pure D_2_O reference samples were measured at 280, 295, and 310 K. For reference, a pure D-Serine solution at 45 mg/mL was also measured on IN5 at one temperature (295 K).

### 2.9 Data reduction and fit algorithms

The Mantid software pacakage [33] was used for the initial reduction of the neutron data recorded at IN16B. The Lamp software package ^1^ provided by the ILL was used for the IN5 data. All fits were carried out using python3 scripts employing the *scipy.optimize.curve_fit* command. Errors in the fit parameters were calculated from the square root of the diagonal of the covariance matrix. The Voigt profiles used to calculate the scattering functions convoluted with the energy resolution functions were obtained from the real part of the Faddeeva function provided by *scipy.special*.

### 2.10 Molecular dynamics simulations

MD simulations were performed using Gromacs 2016.3 [34, 35, 36] with the Amber99SB-ILDN force field.[37] Parameters for the D-Serine ligand were derived from the existing L-Serine parameters. Charges for D-serine were obtained by the RESP approach, as described in [38, 39, 40]. Quantum mechanical calculations prior to RESP calculations were done with TURBOMOLE V7.1 [41] on the Hartree-Fock level using the RI-J approximation[42] and a 6-31G* basis set.[43,44, 45] Two different side-chain conformers of D-Serine were used and charges averaged over these two conformations.

For the pyruvate residue, existing force-field parameters from acetate and the amide carbonyl were used. Charges were calculated as described above using a single conformation.^2^

For analysis and visualisation of MD trajectories we used self-written Python scripts in combination with the modules MDAnalysis,[46] NumPy[47] and Matplotlib.[48]

The ADC-D-Serine complex determined in this work and the ADC-APO structure (1AW8 [19]) were used as the starting structures for the MD simulations. For each simulation an ADC tetramer (apo and D-Serine bound) was placed in a cubic box with periodic boundary conditions (1 nm initial minimum distance of protein to all boundaries). The box was filled with water (ca. 48000 molecules). To neutralize the negative charge of the protein, a number of water molecules were replaced with sodium and chloride ions to reach a concentration of 100 mM of NaCl. For each system MD simulations were prepared at two different temperatures (285 K and 310 K). After an energy minimisation (50000 steps or maximum force *<* 10 kJ mol^−1^ nm^−1^), an NVT equilibration with modified Berendsen thermostat and velocity rescaling [49], and a 0.1 ps timestep (separate heat bath for protein and solvent+ions) was run. This was followed by a NPT equilibration using a Parrinello-Rahman pressure coupling [50, 51] at 1 bar with a compressibility of 4.5×10−5 bar^−1^ and a 2 ps time constant. During both runs, a position restraint potential with a force constant of 1000 kJ mol^−1^ nm^−2^ was added to all protein atoms (including the ligand). All bonds to hydrogen atoms were constrained with the Linear Constrained Solver (LINCS)[52] with an order of 4 and one iteration. MD simulations were run with a time step of 1 fs and the leapfrog integrator. Coordinates were saved every 10 ps. A grid-based neighbor list with a threshold of 1 nm was used and updated every 10 fs. For long-range electrostatic interactions above 1 nm the particle-mesh Ewald method[53, 54] was used with a fourth order interpolation and a maximum spacing for the FFT grid of 1.6 Å. Lennard-Jones interactions were cut-off above 1 nm. A long range dispersion correction for energy and pressure was used to compensate for the Lennard-Jones interaction cut-off.[35] For each of the four MD simulations 250 ns were acquired.

## 3 Results

The crystal structure of Aspartate *α*-decarboxylase in complex with D-Serine was determined to a resolution of 1.9 Å. The structure was refined to final crystallographic R_*work*_ and R_*free*_ values of 16.7% and 18.6% respectively (Table: 1). As in the apo-ADC stucture, the ADC tetramer is formed by a crystallographic two-fold, with two ADC monomers in the asymmetric unit. As in the apo structure, a fraction of mis-processed ADC is present, where the backbone of the zymogen is cleaved, but the *β* subunit has an N-terminal serine instead of a pyruvate. This mis-processed form is present at 40 % occupancy.

**Table 1:** Crystallographic refinement statistics

|  |  |
| --- | --- |
|  | ADC-D-Serine complex |
| Wavelength (Å) | 0.9778 |
| Resolution range (Å) | 46.93 - 1.75 (1.81 - 1.75) |
| Space group | P 61 2 2 |
| Unit cell (Å) | 71.3 71.3 216.6 90 90 120 |
| Total number of reflections | 67840 (6601) |
| Number of unique reflections | 33921 (3301) |
| Multiplicity | 2.0 (2.0) |
| Completeness (%) | 99.99 (99.97) |
| Mean I/sigma(I) | 21.50 (5.25) |
| Wilson B-factor (Å<sup>2</sup>) | 19.07 |
| R-merge | 0.02384 (0.1564) |
| R-meas | 0.03371 (0.2212) |
| R-p.i.m. | 0.02384 (0.1564) |
| CC1/2 | 0.99 (0.93) |
| CC* | 1.00 (0.98) |
| Reflections used in refinement | 33921 (3301) |
| Reflections used for R-free | 1702 (162) |
| R-work | 0.1791 (0.2110) |
| R-free | 0.1937 (0.2266) |
| CC(work) | 0.956 (0.836) |
| CC(free) | 0.957 (0.807) |
| Number of non-hydrogen atoms | 2172 |
| macromolecules | 1950 |
| ligands | 14 |
| solvent | 208 |
| Protein residues | 244 |
| RMS bonds (Å) | 0.014 |
| RMS angles (°) | 1.67 |
| Ramachandran favored (%) | 96.51 |
| Ramachandran allowed (%) | 3.06 |
| Ramachandran outliers (%) | 0.44 |
| Rotamer outliers (%) | 3.5 |
| Clashscore | 5.15 |
| Average B-factor (Å<sup>2</sup>) | 23.17 |
| macromolecules | 21.83 |
| ligands | 27.57 |
| solvent | 35.45 |

Comparison of the ADC-APO structure (1AW8) with the D-serine complex shows that the *α* C-terminal loop opens upon D-Serine binding (Figure 1).

**Figure 1:**
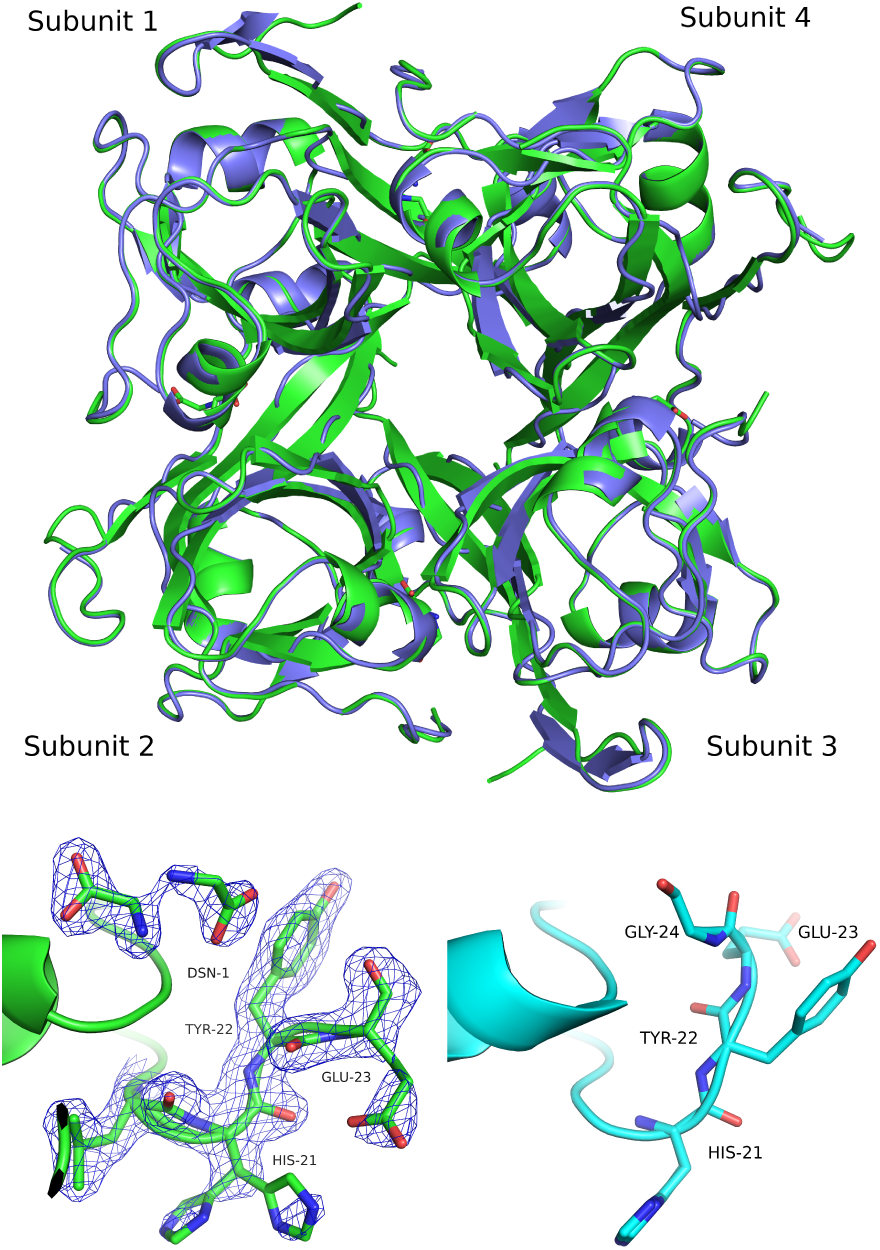
Top: superposition of ADC-APO (1AW8) (blue) and ADC-D-Serine complex structure (green). Bottom: Change in the conformations of the *α* C-terminal loop upon binding of D-Serine in subunit 3 (bottom left) and the corresponding loop conformation in ADC-APO (PDBID 1AW8) (bottom right). 2mF_*o*_-DF_*c*_ electron-density map is displayed at a contour level of 1*σ*.

The D-Serine molecule adopts two conformations in both subunits of the asymmetric unit. One conformer (60% occupancy) forms hydrogen bonds with the mainchain of Ala 75, the sidechains of Arg 54 and Thr 57, and the pyruvoyl carbonyl (Supplementary Figure S1) adopting a similar binding conformation to the native substrate L-aspartate. The second conformer (40% occupancy) forms hydrogen bonds with the sidechains of Lys 9 and Tyr 58 (Supplementary Figure S1) and with the nitrogen atom of the misprocessed *β* N-terminal Ser 25.

In the apo-ADC structure the *α* C-terminal loop (residues 21 to 24) adopts two conformations, whereas in the D-serine complex, the loop adopts a single, open conformation (Figure 1). The degree of “openness” is not the same in the two ADC subunits of the asymmetric unit. There is a significant change in the conformation of the *α* C-terminal loop of chain A (subunit 1) which shows a displacement of 4.3 Å of residue Glu 23 from its position in the apo structure. However, for chain D (subunit 2), the structural change is relatively small with a C_*α*_-C_*α*_ distance of only 0.8 Å for the same residue (Figure: 1), although D-serine is bound in both subunits.

This observation is supported by the MD simulations of apo and D-serine complexed ADC. The conformational change which is associated with the displacement of the C-terminal loop of subunit 3 occurs mainly between HIS 21 and GLY 24. Hence, we monitored the change in the C_*α*_-C_*α*_ distance betweem His21 and Gly24, and between Tyr22 and Gly24. The distance histogram profiles are significantly different for ADC-APO and ADC-LIG. The histogram profile on the top panel shows three peaks for the His21-Gly24 distance in ADC-APO at 8.2, 9.7 and 10.5 Å whereas for the D-serine complex, there are just two peaks at 8.7 and 9.5 Å. Similarly, there is a clear shift in the distance distribution of Tyr22-Gly24 from 7.25 Å in ADC-APO to 5.75 Å in the D-serine complex. This indicates that the motion of the C-terminal loop is relatively more confined in the presence of ligand than in ADC-APO (Figure 2). Further, the changes in the two distances for the individual subunits support the observation from the crystal structure that the influence of binding of D-Serine on the dynamics of the C-terminal loop is neither completely symmetrical nor consistent among the four subunits.

**Figure 2:**
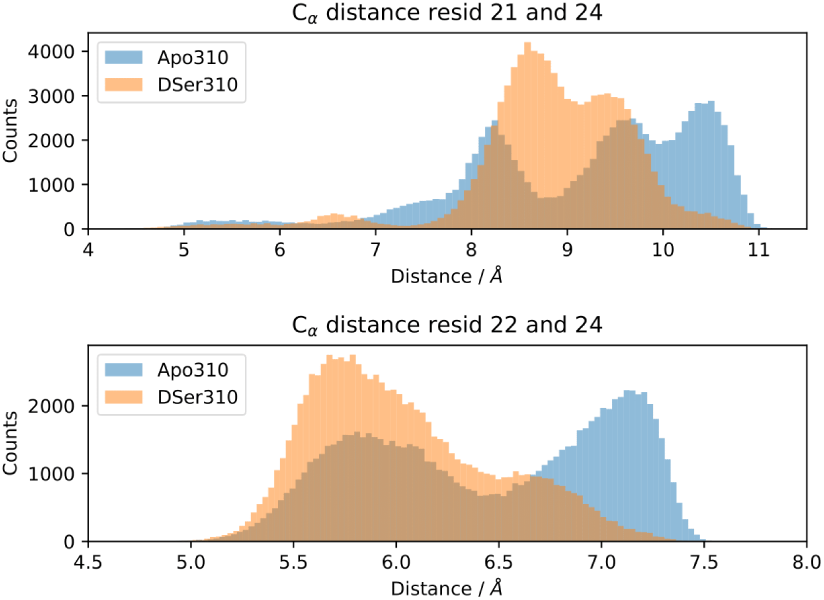
Histograms for the average C_*α*_-C_*α*_ distances for all the four subunits between HIS 21-GLY 24 (top) and between TYR 22-GLY 24 (bottom) for ADC-APO (blue) and ADC-LIG (orange).

The data from the time-of-flight spectrometer IN5 contain information on both quasielastic scattering arising from diffusive motions and inelastic scattering arising from vibrational motions. Here, we present the QENS part only. The reduced QENS data from IN5 were fitted for each momentum transfer *q* independently in a two-step procedure as follows. First, the spectra from the buffered D_2_O solvent were fitted by [55]

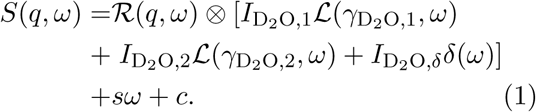

Therein, *ℛ*(*q, ω*) represents the apparent energy resolution function of IN5, which also includes effects from the sample container geometry, and ⊗ the convolution in the energy transfer ħ*ω*. Technically, the convolution *ℛ*⊗ is carried out by modeling *ℛ* as a sum of an arbitrary number of Gaussian functions. Therefore, the observable *S*(*q, ω*) can be fitted by a sum of Voigt functions [12, 56, 57]. *ℒ*(*σ, ·*) represents a Lorentzian function with the width *σ*, and *δ*(*ω*) is the Dirac function describing the elastic scattering arising from the sample container. 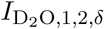 denote scalar *q*-dependent intensity factors, and *s* and *c* are *q*-dependent scalars describing an apparent background arising from the sample, container, and instrument itself.

Second, the QENS spectra from the protein solution samples and pure D-Serine reference sample are fitted by

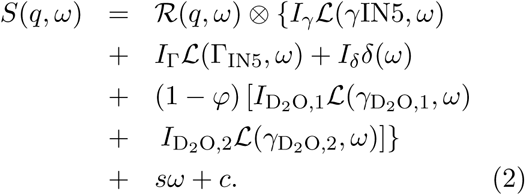

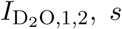, and *c* are fixed from the fit results of the corresponding pure solvent (equation 1), and *φ* is the known protein volume fraction in the sample. The term *I*_*δ*_*δ*(*ω*) accounts for the apparent elastic scattering arising from both the sample container and sample dynamics that are quasi-static on the observation scale of IN5. The fit results for the Lorentzian widths assigned to ADC in equation 2 are depicted in figures 4 and 5.

**Figure 3:**
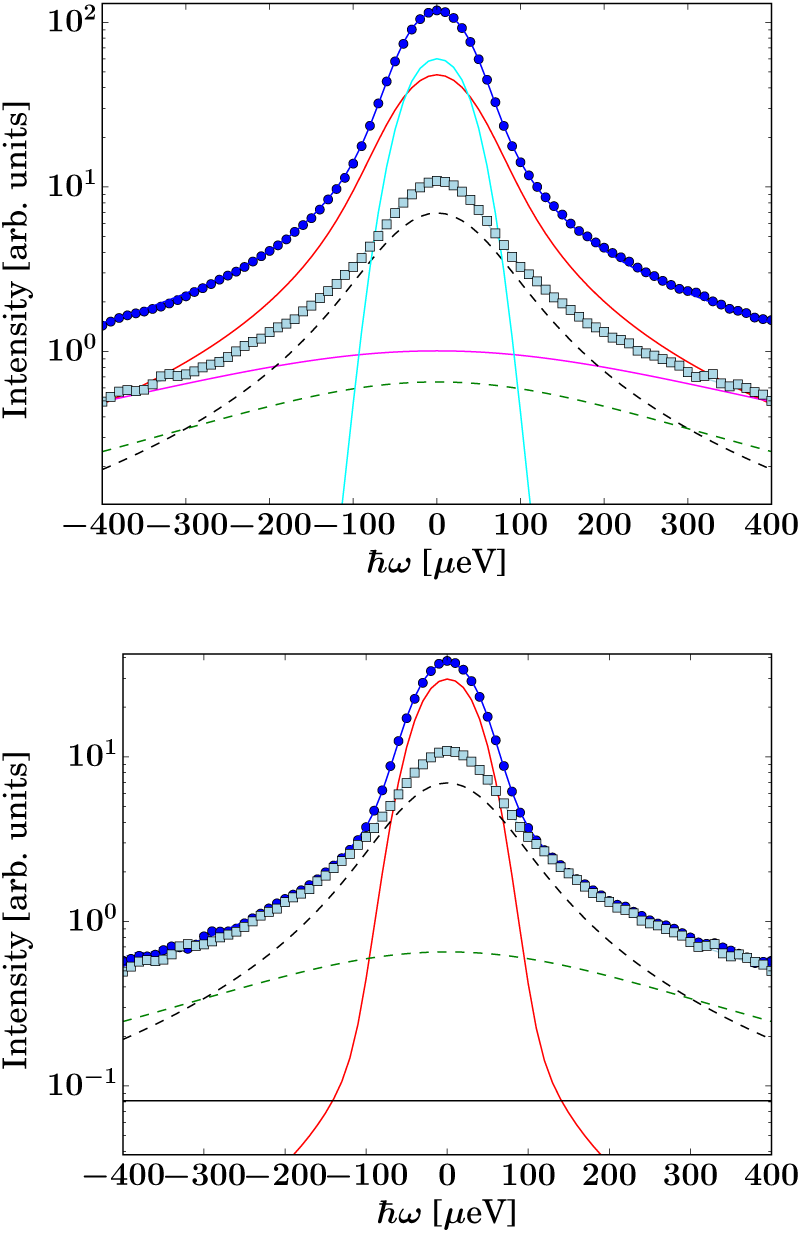
Example spectra recorded on IN5 on ADC-Lig. (top) and ADC-APO (bottom), respectively, (symbols) at *q* = 0.6 Å^−1^ at *T* = 295 K. The dark blue circle symbols represent the protein solution signal, the light blue square symbols the corresponding solvent signal. The lines superimposed on the protein sample spectra represent fits of equation 2. The dashed lines represent the two Lorentzians describing the buffer, equation 1. The other solid lines depict the two Lorentzians associated with the protein and apparent elastic contribution, equation 2. The difference in the spectral intensity between the two samples arises from the D-Serine present in the ADC-Lig sample.

**Figure 4:**
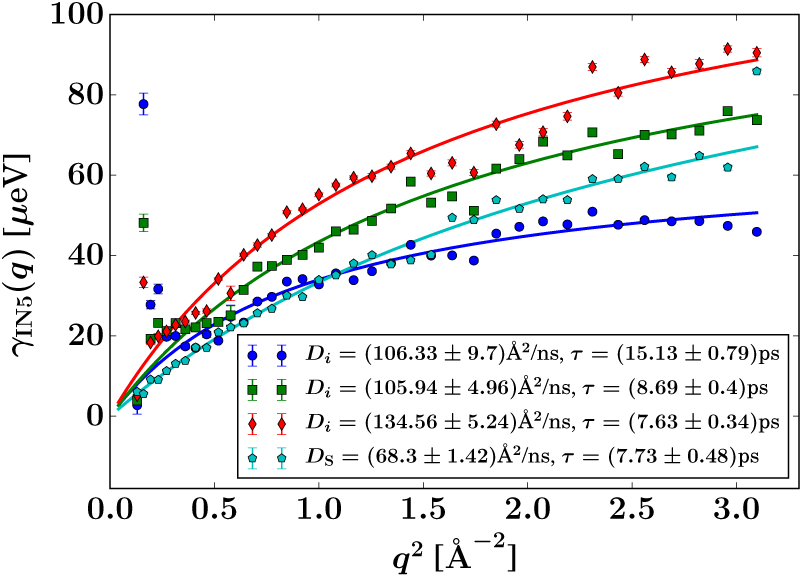
Width *γ*_IN5_ of the Lorentzian accounting for slow internal diffusive motions observed on IN5 (equation 2) for ADC-Lig at different temperatures (circles – 280 K, squares – 295 K, and diamonds – 310 K) as well as for the pure D-Serine reference sample (pentagrams – 295 K), and fits using the jump diffusion model (equation 5).

**Figure 5:**
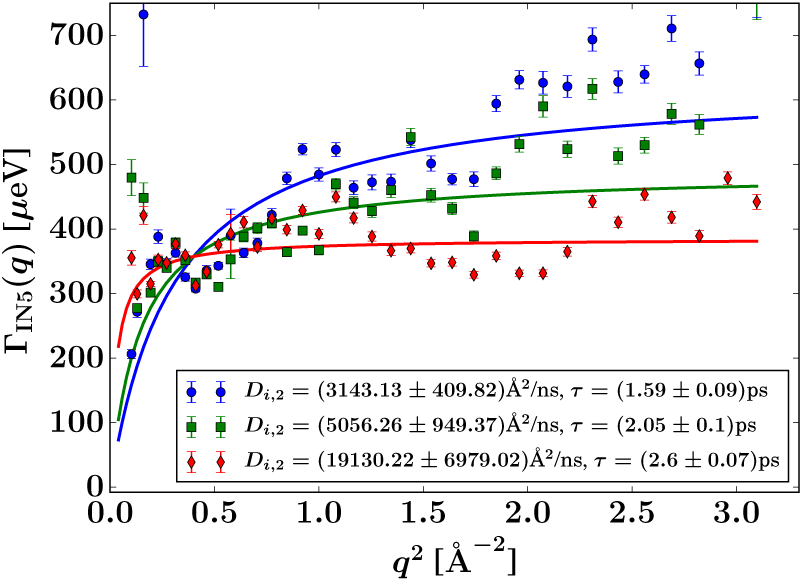
Width Γ_IN5_ of the Lorentzian accounting for fast internal diffusive motions observed on IN5 (equation 2) for ADC-Lig at different temperatures (symbols) and fits using the jump diffusion model (equation 5), the fit parameters being reported in the legend.

In the case of the ADC-APO sample, the amplitude of the Lorentzian associated with the slower part of the internal diffusive dynamics is very small (figure 4). In contrast, this contribution is significant in the pure D-Serine reference sample (figure 4), although with a different linewidth, at the same measured temperature of 295 K. In the ADC-D-Ser samples, the first Lorentzian can thus be attributed to the dynamic contribution of the D-Serine. It should be noted that this Lorentzian with width *γ*_IN5_ in equation 2 accounts for an average over multiple dynamic contributions that cannot be further discerned with the current accuracy of the data and modeling. In the case of the ADC-D-Serine sample, this Lorentzian likely reflects both bound and unbound D-Serine. The observed diffusion coefficient of pure D-Serine is in reasonable agreement with earlier findings [58, 59, 60].

Overall, it appears that few accessible diffusive dynamic contributions on the picosecond time scale are associated with the protein itself, suggesting an overall highly rigid protein consistent with its high content of *β*-sheet (approximately 40%) as determined using the DSSP server [61, 62].

With its high energy resolution, the spectrometer IN16B accesses quasi-elastic scattering containing information on superimposed center-of-mass and internal diffusion. The following model was used for the scattering function observable on IN16B [12, 56, 57]:

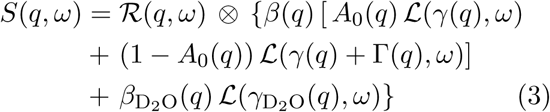

Where, *ℛ* = *ℛ* (*q, ω*) denotes the spectrometer resolution function, and *ℒ* (Δ, *σ*) is a Lorentzian function with the HWHM *σ. β*(*q*), *A*_0_(*q*), *γ*(*q*), and Γ(*q*) are scalar fit parameters. The scalar parameters for the solvent water contribution 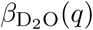 and 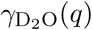 were fixed based on a pure solvent measurement using established protocols [63].

Using a global fit approach of the spectra for all *q* simultaneously using equation 3, a Fickian center-of-mass diffusion of the proteins described by the observable apparent diffusion coefficient *D* was assumed, using the following relation which has been determined for other proteins [56, 57, 63],

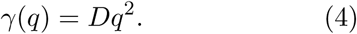

To examine the internal diffusion, a similar global fitting approach was taken, using equation 3, and assuming a jump diffusion model for the internal diffusion within ADC [64]

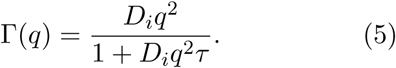

where, *D*_*i*_ is the jump diffusion coefficient and *τ* is the so-called residence time between diffusive jumps. In earlier work [11], this jump diffusion model has been shown to be a reasonable approximation. Importantly, in the fits of the IN16B spectra, the values of *D*_*i*_ and *τ* are fixed based on the results from IN5 for ADC-Lig (figure 4). An example spectrum and fit using equation 3 is shown in figure 6.

**Figure 6:**
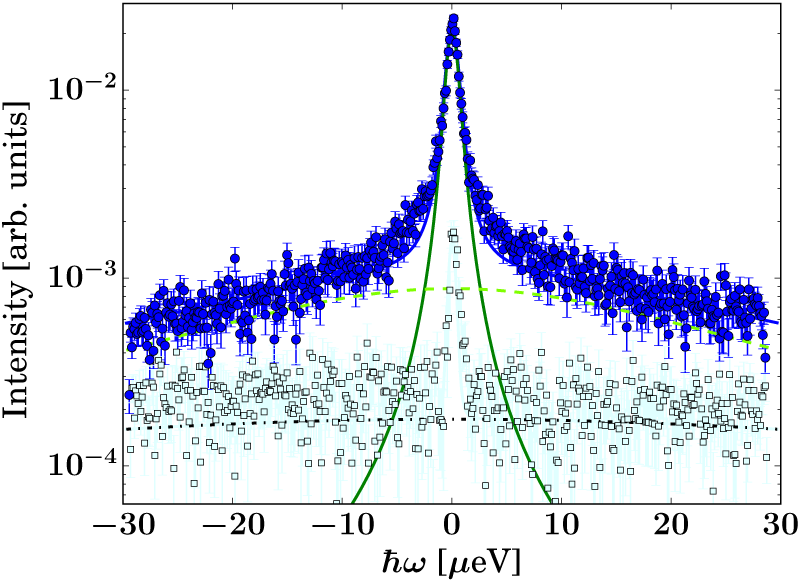
Example spectrum (filled circles) of ADC-Lig recorded on IN16B at *T* = 295 K, *q* = 0.78 Å^−1^ and fit using equation 3 (solid line superimposed on the symbols consisting of the components (other lines) as given by eq. 3). The lower light blue open circles represent the pure solvent signal.

In equation 3, *A*_0_(*q*) can be identified with the elastic incoherent structure factor, EISF [65] (figure 7) as follows:

**Figure 7:**
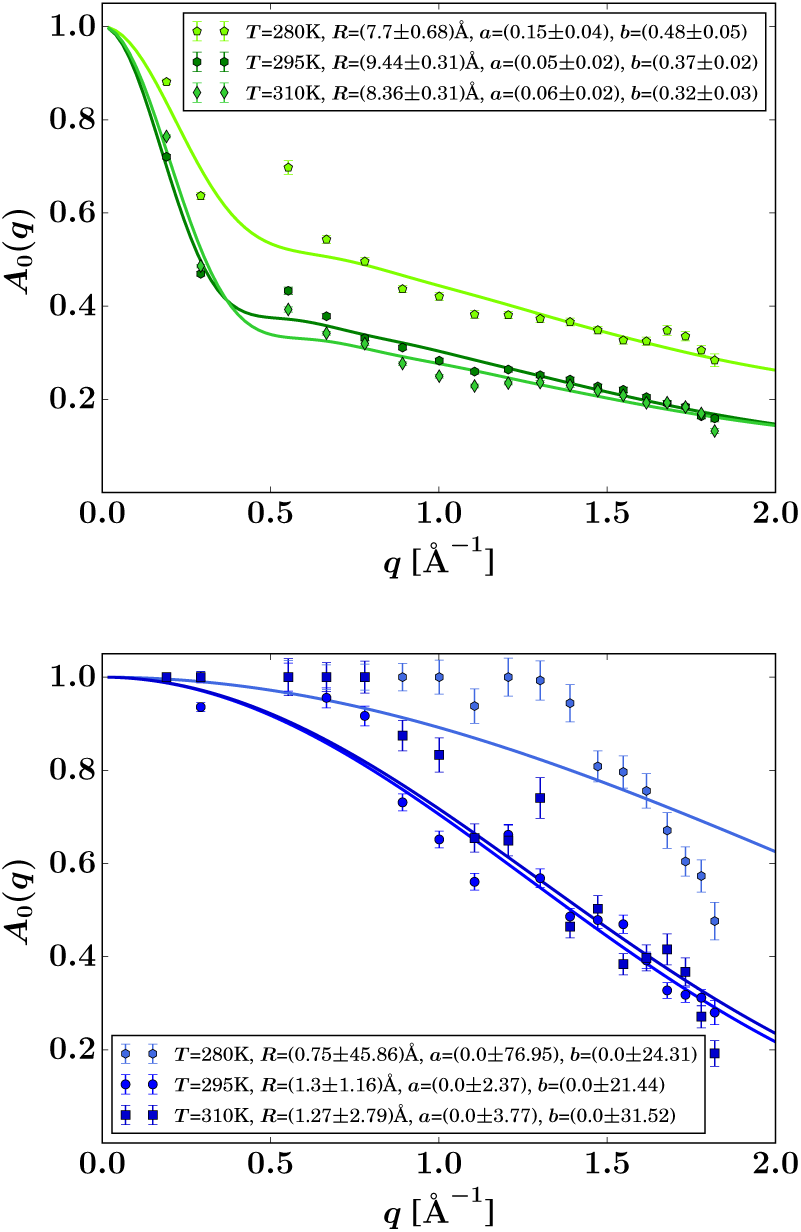
EISF *A*_0_(*q*) (symbols) for ADC-Lig (top) and ADC-APO (bottom) obtained from equation 3 when fixing Γ by using the IN5 result, and fits by equation 6 (solid lines). The resulting fit parameters are given in the legends.

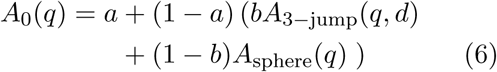

where, *a* is the fraction of scatterers within the protein that are immobile (apart from the protein center-of-mass diffusion) on the observation time or coherence time of the measurement (on the order of a nanosecond for IN16B), and *A*_3−jump_(*q, d*) describes atoms undergoing reorientational jumps between three equivalent sites, which are associated with the methyl groups,

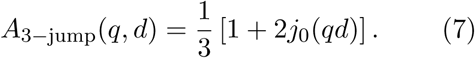

The jump length *d* = 1.715 Å is fixed for these methyl groups [57, 65, 66]. it is further assumed that the protein side chains diffuse on the surface of a sphere with the average radius *R*, such that

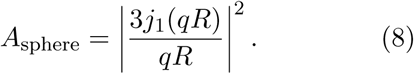

where, *j*_0_ and *j*_1_ are the spherical Bessel functions of zeroth and first order, respectively. *b* denotes the relative weight of the contributions from *A*_3−jump_(*q*) and *A*_sphere_. Results for the EISF *A*_0_(*q*) and fits using equation 6 are reported in figure 7 for both samples at the higher protein concentration *φ* = 0.09.

We note that the fit result for the EISF is strongly dependent on the fixing of the linewidth Γ in equation 3 using the fit results from IN5. In contrast, when not fixing the internal dynamics (not shown), a finite internal linewidth Γ(*q*) in equation 3 can only be seen for the samples with the ligands. For the Apo-samples, Γ(*q*) *→ ∞* in the fits of the IN16B spectra. At the same time, the errors on the internal diffusion fit parameters diverge. This could suggest very fast internal motions beyond the dynamic window accessible by IN16B.

Remarkably, the internal diffusive dynamics seem to change substantially depending on whether or not the ligand is present, although this it should be noted that this change may simply reflect the dynamics of the D-Serine itself. However, independently from the assumptions on the internal dynamics, the fit results for the center-of-mass diffusion coefficient *D* appear rather robust. The systematic error due to assumptions on the internal dynamics is on the order of *±*0.5Å^2^*/ns*, thus, larger than the fit parameter confidence bounds, but significantly smaller than the difference in the diffusion between the too samples (table 3).

**Table 2:** HYDROPRO output for ADC-APO (PDBID 1AW8) and ADC-LIG structures for 280, 295 and 310K temperatures

|  |  | ADC-APO |  |  | ADC-LIG |  |
| --- | --- | --- | --- | --- | --- | --- |
| PDB file (tetramer) | 1AW8 | 1AW8 | 1AW8 | This work | This work | This work |
| Radius of elements in primary model ($\text{\AA}$) | 2.90 | 2.90 | 2.90 | 2.90 | 2.90 | 2.9 |
| Temperature (K) | 280 | 295 | 310 | 280 | 295 | 310 |
| Solvent viscosity (pure D2O) (poise) | 0.01830 | 0.01175 | 0.00830 | 0.01976 | 0.01270 | 0.00896 |
| Molecular weight (Da) | 50391 | 50391 | 50391 | 50811 | 50811 | 50811 |
| Solute partial specific volume ($\text{cm}^3/\text{g}$) | 0.702 | 0.702 | 0.702 | 0.702 | 0.702 | 0.702 |
| Solution density ($\text{g}/\text{cm}^3$) | 1.110 | 1.110 | 1.110 | 1.110 | 1.110 | 1.110 |
| Translational diffusion coefficient ($D_t(\varphi=0)$) ($\text{cm}^2/\text{s}$) | 3.784E-07 | 6.209E-07 | 9.236E-07 | 3.163E-07 | 5.188E-07 | 7.721E-07 |
| Radius of gyration (cm) | 2.257E-07 | 2.257E-07 | 2.257E-07 | 2.446E-07 | 2.446E-07 | 2.446E-07 |
| Volume ($\text{cm}^3$) | 8.235E-20 | 8.235E-20 | 8.235E-20 | 9.476E-20 | 9.476E-20 | 9.476E-20 |
| Rotational diffusion coefficient ($D_r(\varphi=0)$) ($\text{s}^{-1}$) | 3.174E+06 | 5.208E+06 | 7.747E+06 | 2.124E+06 | 3.484E+06 | 5.185E+06 |

**Table 3:** Diffusion coefficients obtained from QENS, DLS and HYDROPRO at 295K in Å^2^/ns. The errors denote 67% confidence bounds on the fits and do not account for systematic errors arising from the choice of the model.

| | QENS $D(\varphi=0.09)$ | DLS $D_t(\varphi \approx 0)$ | HYDROPRO $D_t(\varphi=0)$ |
| --- | --- | --- | --- |
| ADC-APO | $2.82 \pm 0.03$ | $4.30 \pm 0.03$ | 6.21 |
| ADC-LIG | $1.39 \pm 0.02$ | $4.05 \pm 0.02$ | 5.19 |

The global apparent center-of-mass diffusion *D* obtained from fitting equation 3 is expected to follow a Stokes-Einstein temperature dependence

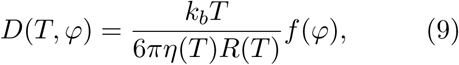

where *η* is the solvent viscosity and *R* the effective protein hydrodynamic radius. *f* (*φ*) is a scalar function of the protein volume fraction *φ* and does not depend on *T* or *R* [32, 56, 67]. Thus, *D* directly reflects the average hydrodynamic size of the diffusing particle, which can be a protein monomer, oligomer, or cluster.

Interestingly, the results in figure 8 indicate that the hydrodynamic size of the ADC tetramer or aggregate depends strongly on whether or not ligand is present. For ADC-APO, a perfect Stokes-Einstein dependence, equation 9 can be observed, as illustrated by the linear dependence on *T*, suggesting that the size of the ADC-APO assembly is constant within the observed temperature range. In contrast, for ADC-Lig, a larger assembly seems to be present, although there is evidence of partial dissociation at higher temperatures. Since the crystal structure indicates that ligand binding does not significantly alter the hydrodynamic size of the ADC tetramer (table 2), the difference between ADC-APO and ADC-Lig samples is best explained by the formation of a higher order protein oliogmer in solution in the presence of D-serine [68, 32].

**Figure 8:**
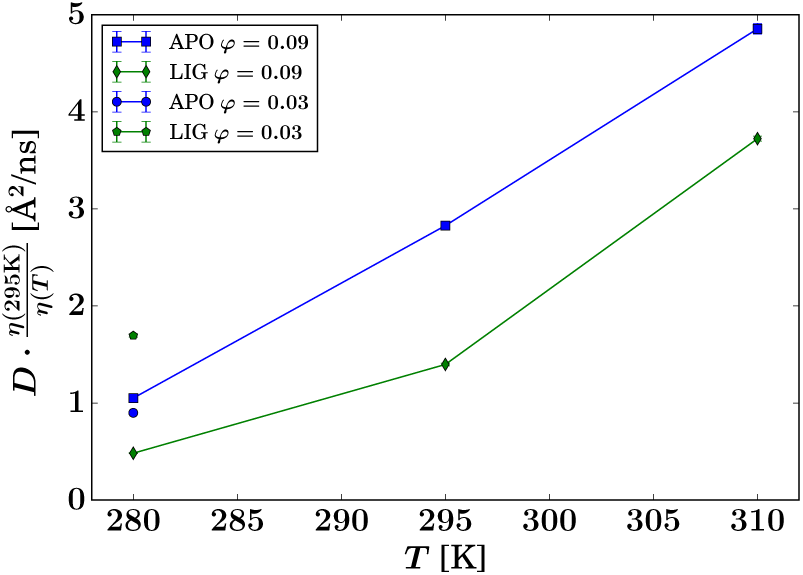
Observable apparent center-of-mass diffusion coefficients *D* (symbols) obtained from the IN16B spectra for all samples and protein volume fractions *φ*, rescaled by the temperature-dependent solvent viscosity *η*(*T*), versus sample temperature *T*. The lines are guides for the eye and do not represent any fit.

In order to further investigate this higher order oligmer formation in the presence of ligand, and ensure that it is not simply a concentration artifact, dynamic light scattering (DLS) measurements were carried out. DLS accesses the collective diffusion of relatively dilute samples as opposed to the self-diffusion in concentrated samples accessed by spatially incoherent QENS. It also observes substantially longer diffusive time scales on the order of milliseconds as opposed to the nanosecond diffusive short-time regime explored by QENS. Due to the low momentum transfers, DLS in general only accesses the translational part *D*_*t*_ of the diffusion coefficient in the case of proteins. Using DLS, a time autocorrelation function was measured over the angular range 30-150 *°* (figure 3). The correlation function for a monodisperse sample is given as

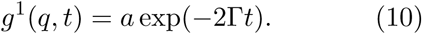

In case of more than one populations of clusters with different diffusion coefficients, the following equation is used as the fit function:

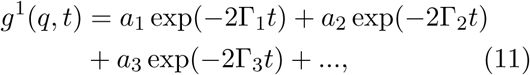

where Γ is the decay rate and *t* the time. The first order autocorrelation function was treated as a monoexponential decay (equation 10) in order to extract the decay rate.

Γ can then be plotted versus q^2^ which follows Fickian diffusion (cf. equation 4),

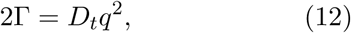

where D_*t*_ is the diffusion coefficient and *q* is the momentum transfer defined as

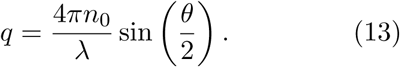

where *θ* is the scattering angle, *n*_0_ the refractive index of the sample, and *λ* the wavelength of the incident beam. Due to the linear relation of Γ and q^2^ (equation 12) a linear fit of the *q*-dependence gives the long-time translational diffusion coefficient *D*_*t*_ (figure 10).

**Figure 9:**
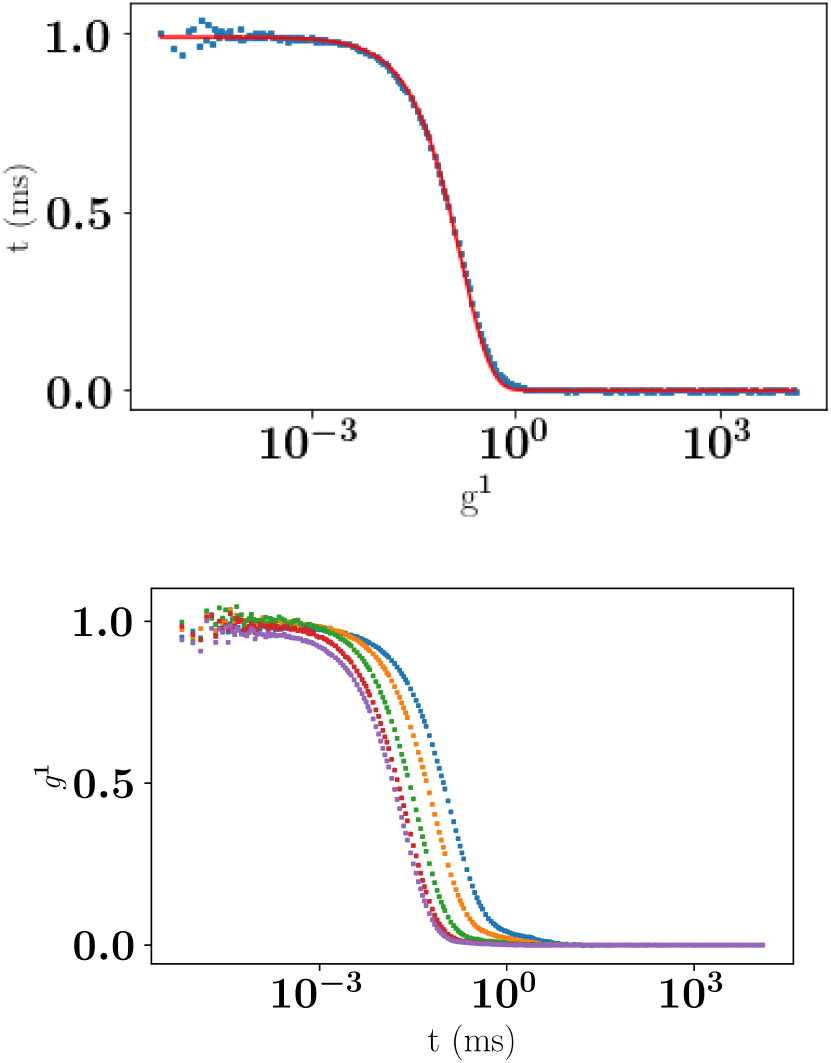
Top: DLS data recorded at Θ = 120*°* and single-exponential fit (equation 10) of the autocorrelation function *g*^−1^ for ADC-LIG at *c*_*p*_ = 15 mg/mL and *T* = 298 K versus correlation time *t*. Bottom: DLS data recorded at 70, 80, 100, 120, 140*°* for the same sample. Symbols represent the data whereas the solid line represents the fit.

**Figure 10:**
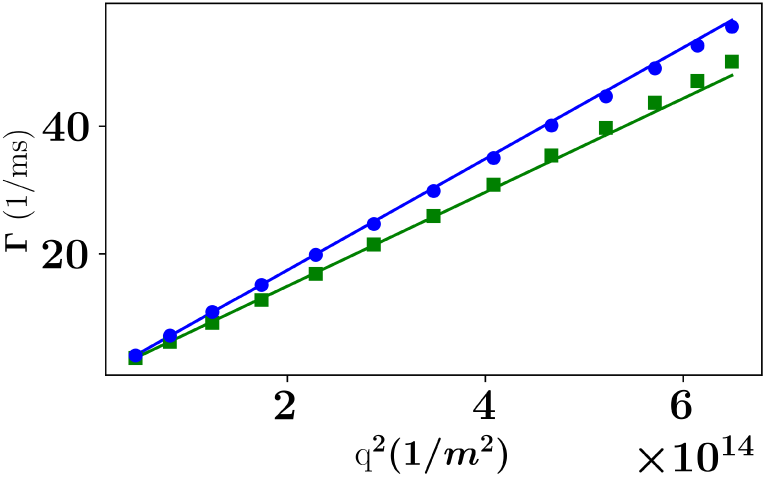
Linear fit of the decay rate obtained from the DLS data for ADC-LIG (squares) and ADC-APO (circles) at 15 mg/mL

By fitting the diffusion coefficients for all five measured concentrations for both the samples ADC-APO and ADC-LIG, average diffusion coefficients were obtained for for both samples at the low-concentration limit (figure 11, symbols at *φ*_*t*_ *≈* 0). These diffusion coefficients of 4.05 *±* 0.02 and 4.30 *±* 0.03 Å^2^/ns for ADC-LIG and ADC-APO, respectively indicate that the two samples behave significantly differently and that the nearly dilute-limit (*φ*_*t*_ *≈* 0) diffusion coefficient is higher for ADC-APO than that for ADC-LIG. This result follows the same trend observed in the QENS results (symbols at *φ*_*t*_ *>>* 0 in figure 11), indicating that the presence of ligand does indeed drive formation of a higher order oligomer.

**Figure 11:**
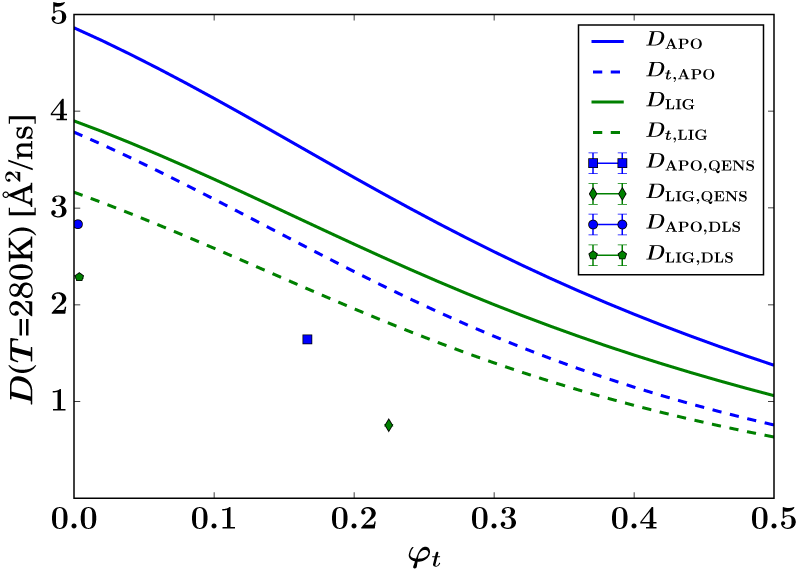
Summary of the QENS and DLS results for the global center-of-mass diffusion coefficient *D* of the proteins versus the theoretical volume fraction *φ*_*t*_ of effective hard spheres, at *T* = 280 K. For DLS, the symbols (at *φ ≈* 0) represent the translational diffusion *D*_*t*_ probed by this method. For QENS, the symbols represent the apparent short-time self-diffusion consisting of contributions from both rotations and translations. The solid lines report the theoretical apparent diffusion of effective spheres with the hydrodynamic size of ADC-APO and ADC-Lig tetramers, respectively. The dashed lines represent the corresponding translational diffusion (cf. legend).

For comparison with the experimental data, the theoretical diffusion coefficients at infinite dilution for the apo-ADC and D-serine complex were calculated using HYDROPRO [26] (Table 2). In general, viscosity depends on both the ionic concentration as well as on the temperature [69, 70, 71]. Changes in viscosity upon addition of L-Serine have been studied previously [69, 72, 70, 71] and the change in viscosity is approximately 5-8% when the serine concentration is increased from 0 to 3.6 M [71]. To account for this increase in viscosity in the calculation of the theoretical diffusion coefficient calculated here, we increased the HYDROPRO input describing the viscosity for deuterium oxide (D_2_O) by 8% and calculated the theoretical diffusion coefficient (Table 2). The translational diffusion coefficients calculated from HYDROPRO for ADC-APO and ADC-LIG are 3.784 and 3.416Å^2^/ns at 280K and 6.206 and 5.623Å^2^/ns at 295K, cf. table 2.

Surprisingly, the lower experimental values from both DLS and QENS compared to the respective theoretical expectations for ADC tetramers indicates that the hydrodynamic size of the experimentally observed diffusing objects is larger than the crystallographic tetramer for both apo-ADC and the D-serine complex. This observation, suggests ADC forms higher order oligomers in solution both with and without D-Serine, although in the prescence of ligand these oligomers are larger.

## 4 Discussion

Previous work on diffusive dynamics in protein solution has focused on standard samples that are commercially available. Here, for the first time, the affect of ligand binding on the diffusive dynamics of a recombinantly expressed non-standard protein sample is studied, combining results from X-ray crystallography, quasi-elastic neutron scattering, dynamic light scattering, and MD simulations. A key challenge for this type of study is discerning the protein internal diffusive dynamics from those of the ligand, as in order ensure to saturation of the protein binding sites a large excess of the ligand must be present in solution. Therefore, although the elastic incoherent structure factor seems to undergo a qualitative change upon ligand binding, this apparent change still has to be determined with higher accuracy. Nevertheless, by combining the QENS data with information from the protein structures, HYDROPRO calculations based on these structures, calculations of the radial hydrogen density distribution functions based on the structures, and theoretical predictions of the short-time self-diffusion of colloidal hard spheres an interpretation of the measured center-of-mass diffusion of the ADC protein in aqueous suspension in the presence and absence of the D-Serine ligand is possible.

The diffusion coefficients obtained from QENS and DLS follow the same trend as the those predicted from calculations, but differ quantitatively. The calculations were made based on the assumption of that the ADC tetramer is the protein assembly present in solution. However, the observed deviation provide strong evidence that larger objects than the ADC tetramers determine the diffusion. Further work will be needed to determine how stable these larger assemblies are, as the time-scales accessed in this study are milli-second and shorter, and how relevant the oligomers are to the biological function of this enzyme. Gel filtration studies show that the dominant species in solution is the ADC tetramer [73], suggesting the higher order species formed here are only transient.

It is clear that ligand binding affects both the nature of these transient oligomers, as well as the internal protein dynamics of ADC. In general, ADC appears rather stiff on the pico-to nanosecond time scale, consistent with the crystallogaphic data (resolution and Wilson B factor). Binding of D-Serine to ADC causes a change in the conformation of the C-terminal loop (Figure: 1), that is observed in the crystal structure, MD simulations and QENS data. Other studies have suggested that the more dynamical a region is in the protein, the more influence it has on the propensity of the protein to aggregate as a result of unfavorable entropic terms [74]. It is therefore perhaps not surprising that small structural changes in highly dynamic regions of the protein, such as the C-terminal loop in ADC, can potentially cause larger changes in the aggregation dynamics of the protein.

## Acknowledgements

TR acknowledges a PhD studentship jointly funded by the ILL and the Universität Hamburg (Federal Excellence Cluster Hamburg Centre for Ultrafast Imaging EXC 1074). We thank the RRZ of the Universität Hamburg for access to their HPC cluster (“Hummel”) for MD simulations and Partnership for Structural Biology (PSB), the Partnership for Soft Condensed Matter (PSCM) in Grenoble for access to their facilities, including notably the DLS. We are grateful to Michael Marek Koza (ILL) for help on IN5 and Ralf Schweins (ILL) for help with the DLS apparatus. BAY acknowledges the Wellcome Trust for supporting this work [110296/B/15/Z].

## Data accessibility

The neutron data are permanently curated by the ILL and accessible under DOI 10.5291/ILL-DATA.8-05-428 [28] and 10.5291/ILL-DATA.8-05-431 [29].

## Supporting Information

### A Amber99SB-ILDN force-field parameters

All parameters given below are in the Gromacs format. We followed the Gromacs manual for adding the residues: http://www.gromacs.org/Documentation/How-tos/Adding_a_Residue_to_a_Force_Field

#### A.1 Atom types, charges and bonded interactions

The lines below were added to aminoacids.rtp in the force field folder. Atom names in the pdb need to be adjusted accordingly before processing.

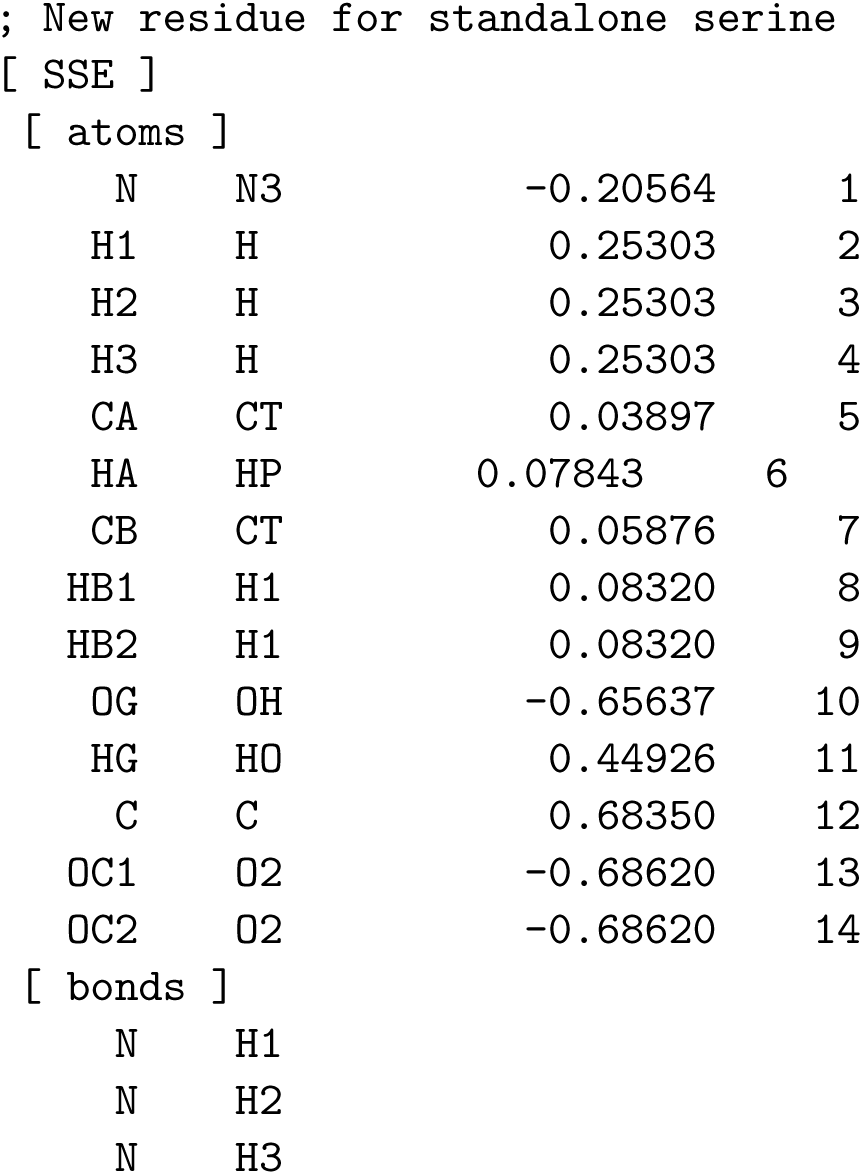
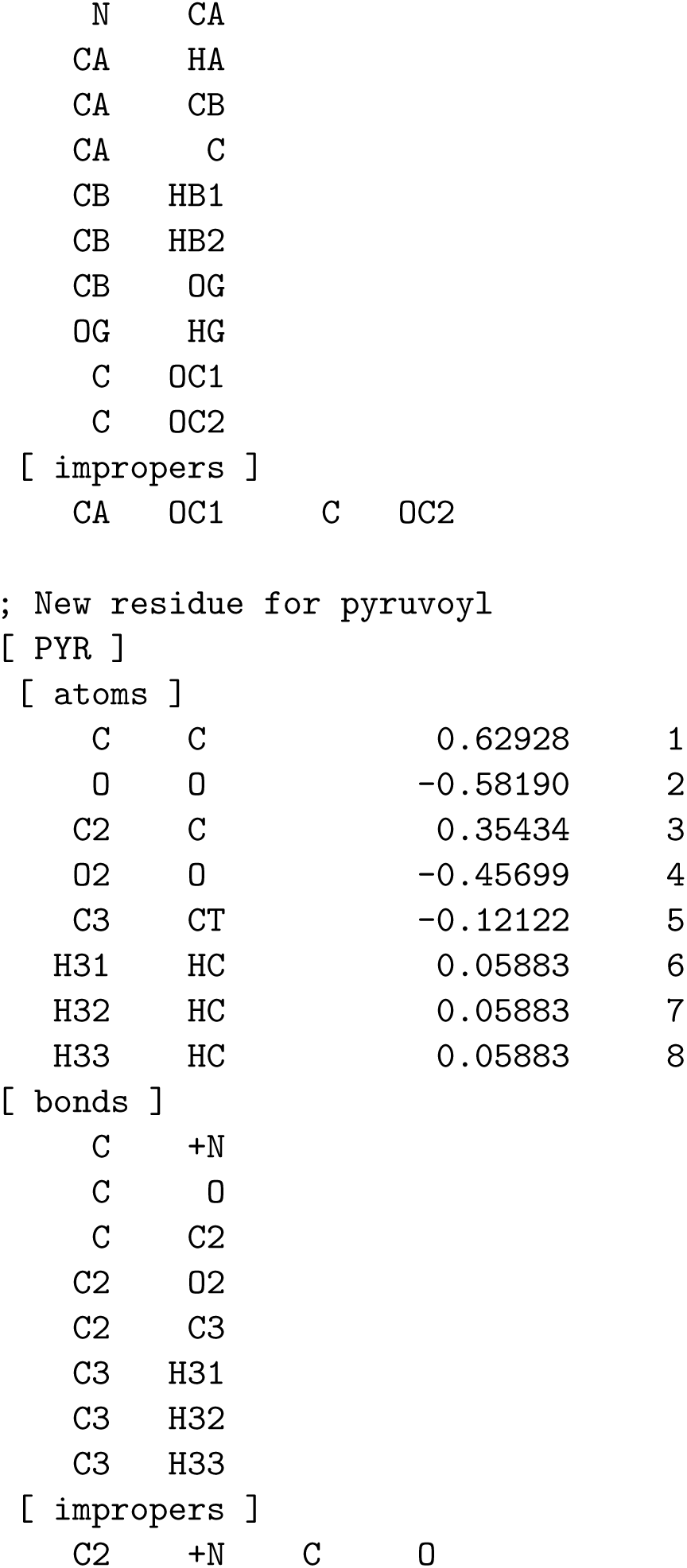

|  |  |  |  |
| --- | --- | --- | --- |
| N | N3 | -0.20564 | 1 |
| H1 | H | 0.25303 | 2 |
| H2 | H | 0.25303 | 3 |
| H3 | H | 0.25303 | 4 |
| CA | CT | 0.03897 | 5 |
| HA | HP | 0.07843 | 6 |
| CB | CT | 0.05876 | 7 |
| HB1 | H1 | 0.08320 | 8 |
| HB2 | H1 | 0.08320 | 9 |
| OG | OH | -0.65637 | 10 |
| HG | HO | 0.44926 | 11 |
| C | C | 0.68350 | 12 |
| OC1 | O2 | -0.68620 | 13 |
| OC2 | O2 | -0.68620 | 14 |

[ bonds ]
|  |  |
| --- | --- |
| N | H1 |
| N | H2 |
| N | H3 |
| N | CA |
| CA | HA |
| CA | CB |
| CA | C |
| CB | HB1 |
| CB | HB2 |
| CB | OG |
| OG | HG |
| C | OC1 |
| C | OC2 |
| [ impropers ] |  |  |  |
| CA | OC1 | C | OC2 |
| C | C | 0.62928 | 1 |
| O | O | -0.58190 | 2 |
| C2 | C | 0.35434 | 3 |
| O2 | O | -0.45699 | 4 |
| C3 | CT | -0.12122 | 5 |
| H31 | HC | 0.05883 | 6 |
| H32 | HC | 0.05883 | 7 |
| H33 | HC | 0.05883 | 8 |

[ bonds ]
|  |  |
| --- | --- |
| C | +N |
| C | O |
| C | C2 |
| C2 | O2 |
| C2 | C3 |
| C3 | H31 |
| C3 | H32 |
| C3 | H33 |
| [ impropers ] |  |  |  |
| C2 | +N | C | O |

##### A.1.1 Addition of hydrogens

This file is necessary, if hydrogens have to be added to the pdb structure. Please add the following lines to aminoacids.hdb.

|  |  |  |  |  |  |
| --- | --- | --- | --- | --- | --- |
| SLE | 6 |  |  |  |  |
| 3 | 4 | H | N | CA | CB |
| 1 | 5 | HA | CA | N | CB |
| 2 | 6 | HB | CB | CA | CG |
| 1 | 5 | HG | CG | CB | CD1 |
| 3 | 4 | HD1 | CD1 | CG | CB |
| 3 | 4 | HD2 | CD2 | CG | CB |
| PYR | 1 |  |  |  |  |
| 3 | 4 | H3 | C3 | C2 | O2 |

##### A.1.2 Translation of atom names in pdb to force-field names

Please add this to the file aminoacids.arn.

|  |  |  |
| --- | --- | --- |
| SSE | O | OC2 |
| SSE | OXT | OC1 |

**Figure S1:**
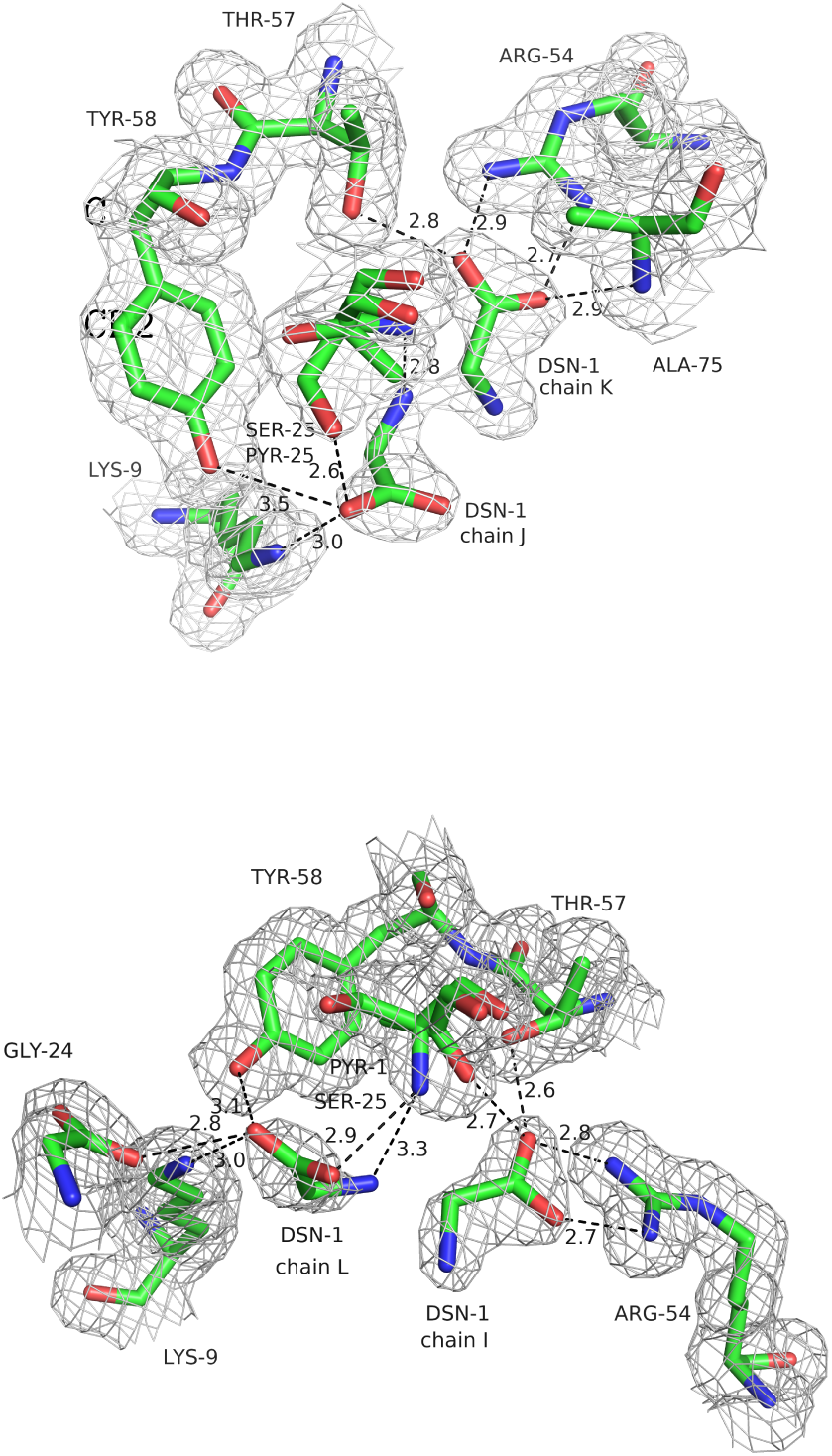
Hydrogen bonding network between the alternate conformations in subunit 1 (top) and subunit 3 (bottom) from the crystal structure of ADC-D-Serine complex showing the slightly different interaction patterns.

**Figure S2:**
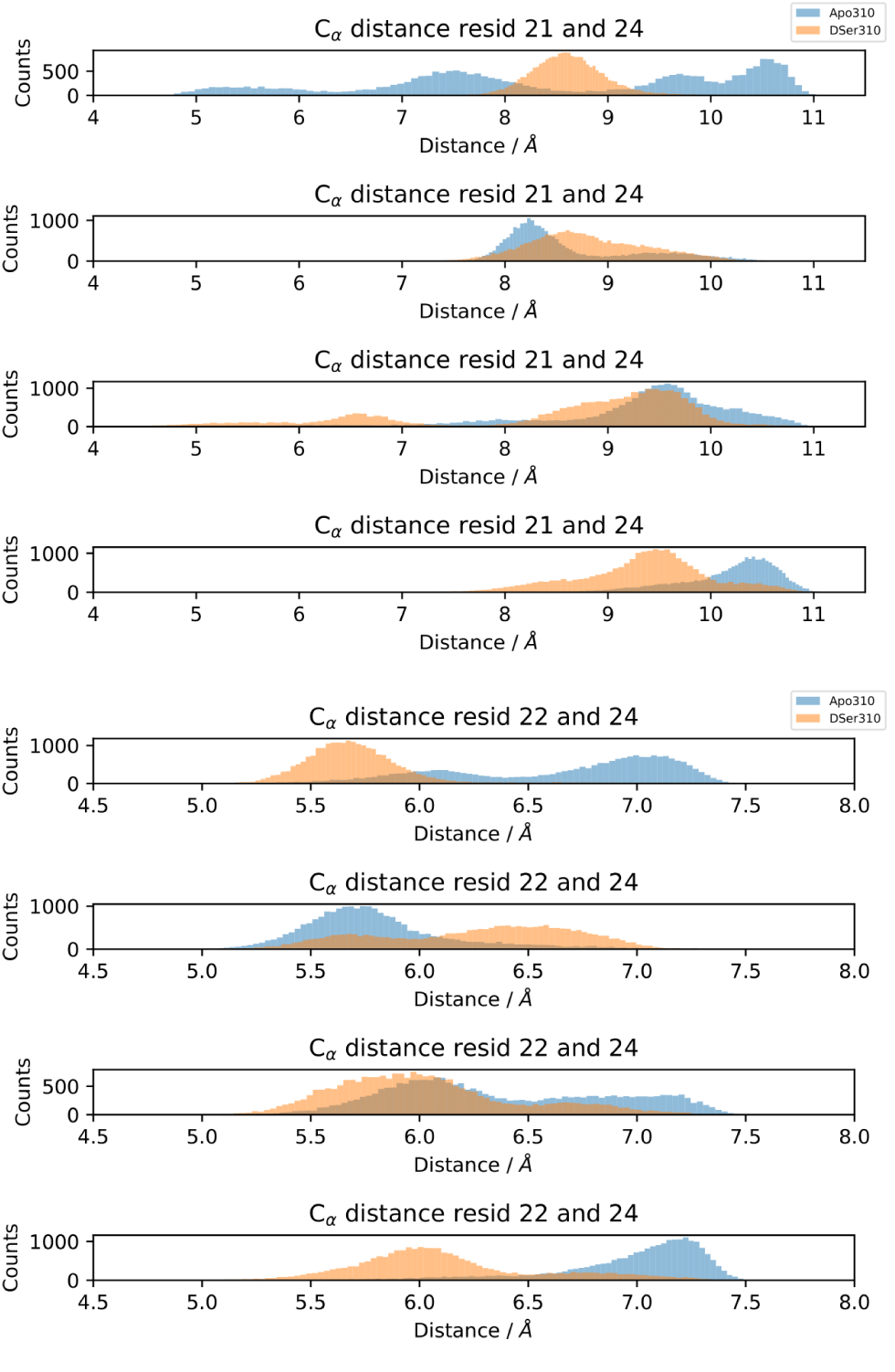
Histograms for the average C_*α*_-C_*α*_ distances for the individual subunits between HIS 21-GLY 24 (top) and between TYR 22-GLY 24 (bottom) for ADC-APO (blue) and ADC-LIG (orange).

## Footnotes

1 For lamp software, please refer to: https://www.ill.eu/users/support-labs-infrastructure/software-scientific-tools/lamp/

2 Force-field parameters for D-Serine and the pyruvate residue are included as supporting information.

